# Mechanical problem solving in mice

**DOI:** 10.1101/2024.07.29.605658

**Authors:** Marcus N. Boon, Niek Andresen, Sole Traverso, Sophia Meier, Friedrich Schuessler, Olaf Hellwich, Lars Lewejohann, Christa Thöne-Reineke, Henning Sprekeler, Katharina Hohlbaum

## Abstract

Behavioral learning is a complex phenomenon that involves cognitive, perceptual, motor and emotional contributions. Yet, most animal studies focus on reductionist tasks that aim to isolate one of these aspects [1, 2, 3, 4]. Here, we use a lockbox — a complex, multi-step mechanical puzzle — to study learning dynamics in freely behaving mice. The mice engaged spontaneously with the task and learned to solve it within just a few trials. To dissect different contributions to this rapid form of learning, we combined deep learning-based behavioral tracking in a multi-camera setup with probabilistic inference and computational modeling. We find that the learning progress of the mice was initially dominated by the acquisition of motor skills, i.e., the increased ability to manipulate the individual mechanisms, while a cognitive strategy for the task sequence emerged only later. The lockbox paradigm may hence offer a promising framework for studying the interaction between low-level motor learning and high-level decision-making strategies in a single, ethologically relevant task.

## Introduction

Mechanical problem-solving [5, 6, 7, 8, 9, 10] and tool use [11, 12, 13, 14] have historically been used as indicators of animal intelligence, and have been observed in a variety of different animals. These complex behaviors, typically seen in species with large brains or high neural densities, suggest advanced reasoning, planning, and causal understanding [15, 16, 17]. However, studying such highly intelligent species, such as corvids, cetaceans, elephants, great apes, and cephalopods, presents significant challenges. Their long lifespans hinder large-scale behavioral studies, while invasive neural recordings and precise genetic manipulations, crucial for understanding the neural basis of behavior, are technically challenging and raise ethical concerns.

For mice, many of the technical challenges have been largely overcome. However, their cognitive abilities are typically studied using relatively simple decision-making paradigms, and even those often require weeks of training to achieve modest performance levels [18, 19]. While simple task designs are instrumental for our understanding of the neural basis of cognition, they offer a limited view of the animals’ natural problem-solving abilities [20, 21, 22, 23, 24] and may underestimate their cognitive potential [25].

To close this gap, we adapted a reemerging sequential decision-making paradigm, mechanical lock-boxes, to mice, to investigate if they can engage in complex mechanical problem solving akin to tasks used to assess intelligence in other species. Lockboxes were previously employed as environmental enrichment for mice to enhance housing conditions and mitigate boredom [26]. We find that mice can learn to solve lockboxes reliably within just a few trials. Combining deep learning-based behavioral tracking and computational modeling, we decompose the learning process and show that it is initially dominated by the acquisition of motor skills, while hallmarks of a cognitive solution strategy emerge only later.

### Mice learn the lockbox in a few trials

We developed a lockbox consisting of a combination of four interlocked mechanical problems, or “mechanisms”, which must be solved sequentially (Figure 1a). The mechanisms are best understood in reverse order: The mouse can collect a reward upon opening a sliding door, but the door is blocked by a ball that cannot be removed before a stick is pulled out, which in turn cannot be pulled out before a lever is raised. All four mechanisms are designed to be easily movable by the mice, while being difficult enough not to be solved by accident. The mechanical interconnections between the mechanisms are visible to the mice and perceivable through tactile perception, allowing them to observe the effects of their actions. In principle, this allows the mice to infer which mechanism needs to be solved next. The animals can move freely and decide to engage or disengage with the task at any moment (Figure 1b). We often observed anticipatory queuing of the mice at the entrance to the task cage (for exemplary video footage of this behavior see here^1^).

**Figure 1:**
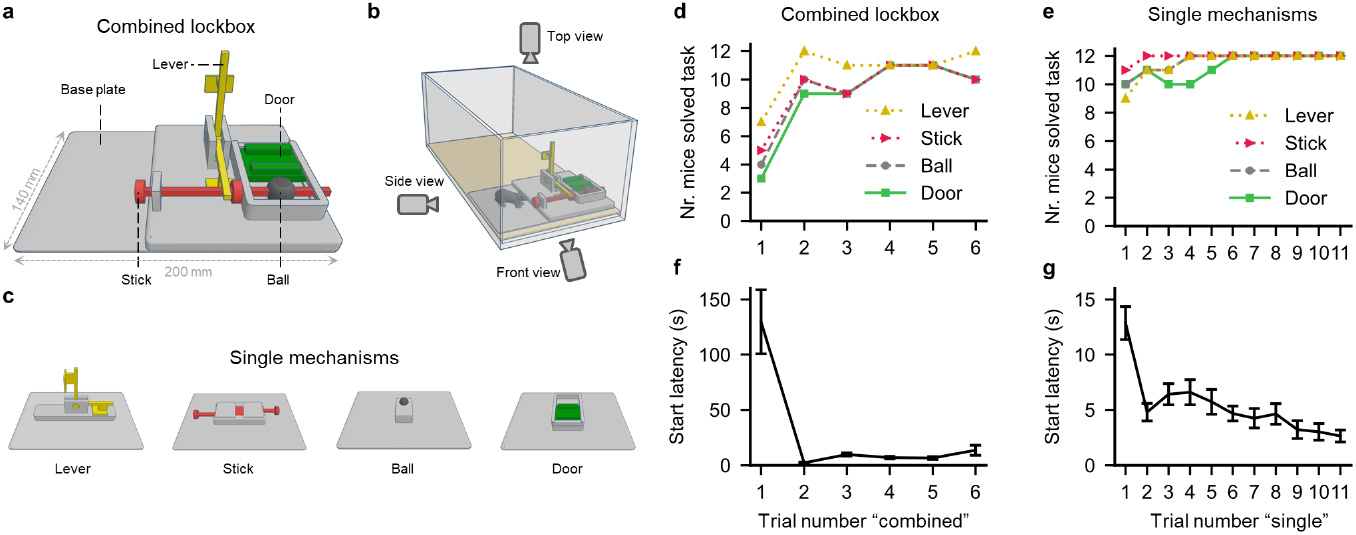
Mice learn the lockbox task rapidly. **a**, 3D model of the combined lockbox. The yellow lever blocks the red stick, which blocks the gray ball, which in turn blocks the green door concealing a food reward underneath. **b**, Schematic depiction of the cage setup. To maintain clarity, the cage grid for food and water, as well as the connection to the second cage, have been omitted from this depiction. The lockbox is placed within a cage and recorded from three perspectives. **c**, 3D models of the single mechanisms in isolation. Each mechanism reveals the reward after a single successful action. In this depiction, the colors of the mechanisms were adapted to match the combined lockbox, but they were yellow in the original prints. **d, e**, Learning curves for the combined lockbox trials and single mechanism trials, respectively. **f, g**, Average values for the first time during each trial that the mice interact with a lockbox mechanism. Error bars represent the standard error of the mean (Fig. f: n = 12, Fig. g: n = [46, 44, 48, 42, 48, 40, 40, 33, 36, 48, 48] for trials 1−11).

We used a three-stage curriculum to train a cohort of young adult female C57BL/6JCrl mice (n = 12). In the first trial, each mouse was exposed to the combined lockbox problem for a maximum duration of 30 minutes. In the subsequent single mechanism training phase (Figure 1c), the mice performed 11 trials in which all individual mechanisms were presented alone for at most 15 minutes each (in randomized order, rewarded per mechanism). As a preference test, all four single mechanisms were then shown simultaneously on two consecutive days (2 trials/day, data not shown here). Finally, the mice were given five trials on the combined lockbox for a maximum duration of 30 minutes on subsequent days. In each trial, if the mouse did not open the lockbox within the specified time, the trial was deemed unsuccessful. If the mouse succeeded in solving the task within the specified time, the trial ended once the mouse reached the reward. For more details on the training paradigm, see Methods and [26].

All mice solved the combined lockbox task at least once within very few exposures. When the mice encountered the task for the first time, only three out of twelve solved it (Figure 1d). After the single mechanism training phase (Figure 1e), there was a sharp rise in the fraction of successful animals (9/12), and this fraction continued to rise slowly across trials except for a decrease by one count in the last trial. The conditional probability that a mouse solves the puzzle, given that it solved it in the immediately preceding trial, was high (*P* (solve_*t*_|solve_*t−*1_) = 0.93). Specifically, 9 out of 12 mice consistently solved the puzzle after their first success; two mice failed only once afterwards, and one mouse increasingly disengaged from the task after succeeding once in the second trial (see Suppl. Inf.).

### Three contributions to learning

We hypothesized that the rapid learning of the task could be shaped by three different contributions: Habituation to the task, motor learning, and task understanding. Here, we distinguish between motor learning as the improvement in the ability to manipulate the individual mechanisms, and task understanding as the acquisition of a sequential solution strategy for the task.

At the onset of the experiment, the lockboxes and the task are novel for the mice. To study the effect of habituation, we analyzed the start latency, defined as the interval between entering the cage and engaging with the lockbox (Figure 1f, g). The start latency was high in the first combined trial (129.8 s *±* 28.9 s, mean *±* SEM, n = 12), but showed a sharp decrease in the second trial (2.2 s *±* 0.72 s) and stayed low in the remaining trials. The mice also displayed a reduction in latency during the single mechanism trials, suggesting that task habituation contributes to the substantial performance increase in the second combined lockbox trial.

To delineate the roles of motor skill acquisition and the development of a cognitive solution strategy, we performed video recordings of the entire training process (*>* 110 h) and implemented an automated video analysis pipeline (Figure 2a). We recorded the behavior from three perspectives and tracked key body parts of the mouse using DeepLabCut (DLC, [27, 28]). The keypoint positions in the three perspectives were combined into a 3D reconstruction using standard triangulation methods and nonlinear temporal filtering with a skeletal model of the mouse (see Methods). From the resulting tracked positions, we classified for each video frame which part of the lockbox the mouse interacted with. We benchmarked the results of the tracking pipeline using manual annotations by two independent human annotators for two videos (Figure 2a, panel (3)). The automated and manual annotations are highly correlated (see Suppl. Inf.). A more extensive comparison can be found in our dataset publication of the experiments [29].

**Figure 2:**
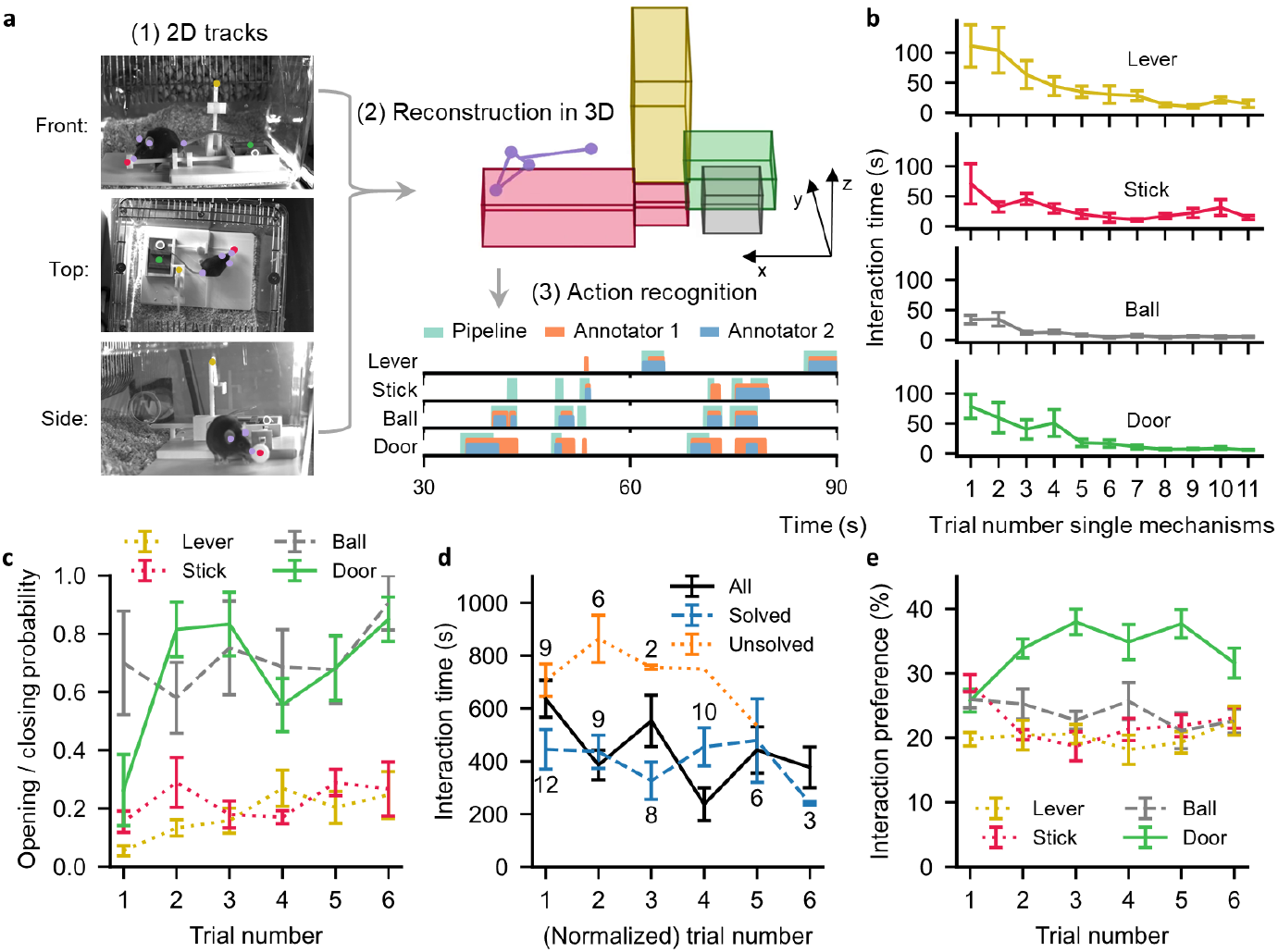
Mice become increasingly proficient in opening the mechanisms. **a**, Schematic of the video analysis pipeline. (1) 2D tracks are extracted from the videos (colored dots). (2) The scene is then reconstructed in 3D and keypoints are used to detect interactions between the mouse (purple) and the lockbox mechanisms (colored blocks). (3) Comparison of pipeline-detected interactions (teal) with two human annotators (orange and blue). **b**, Average interaction time spent on the individual mechanisms during the single mechanism training phase. **c**, Probabilities of mice successfully opening/closing a combined lockbox mechanism during an interaction, given that the mechanism can be opened/closed. **d**, Average interaction time for the combined lockbox for all mice (black), for the trials in which the lockbox was solved (blue), and for the trials where the lockbox was not solved within the specified trial duration (orange). For the latter two, trial numbers are normalized for each mouse such that the first time that a mouse succeeded (blue) or failed (orange) in solving the lockbox is the first data point. The numbers above and below the blue and orange curves represent the number of trials that comprise the data points. **e**, Average fraction of time spent at each mechanism over trials (solved and unsolved). All error bars represent the standard error of the mean, where n = [12, 12, 10, 11, 11, 12] for all combined trial data (c − e). Some trials have fewer data points due to errors in the recording process. For the n value for single trial mechanisms, see Methods.

To analyze the role of motor learning, we first focused on the single mechanism trials. Most mice were able to solve all single mechanisms in the first trial, and performance further increased to 100% for all mice by trial 6 (Figure 1e). The time the mice interacted with the individual mechanisms until they were solved declined by a factor *>* 5 over the course of the 11 trials (Figure 2b, duration trial #1 / #11 for lever: 111 s / 15 s, stick: 71 s / 15 s, ball: 34 s / 5 s, door: 79 s / 6 s). In the subsequent trials with the combined lockbox, the probability of opening or closing a mechanism during an interaction also increases across trials (Figure 2c). Hence, the mice became increasingly proficient at manipulating the individual mechanisms.

For the combined lockbox, we also observe a significant decrease in interaction time across trials (Figure 2d, black curve, p = 0.033 for nonzero correlation, Pearson r = -0.26). However, when we split the trials into success and failure trials, we found no correlation between the interaction time and the number of successes or failures (blue and orange curves, p = 0.62 and p = 0.80, respectively). Thus, the observed decline in interaction time across trials may be a consequence of the increasing ability of the mice to open the full lockbox and thereby end the trial early.

### A strategy emerges for late task stages

To study whether the mice acquired a cognitive strategy for the task, we first quantified how they distributed their time during full lockbox trials. While the overall preferences of the mice remain roughly constant, we observed an increase in the relative time spent with the door mechanism (Figure 2e, p = 0.0498 for nonzero correlation, Pearson r = 0.24), indicating that they learned the reward location. For a more detailed picture, we used a statistical model to infer which of two predefined behavioral strategies best fits the mouse behavior (Figure 3a). The “random” strategy assumes that the mice interacted with the lockbox mechanisms independently of the state of the lockbox, but with mechanism preferences matched to mouse interaction data. The “smart” strategy assumes that the mice preferentially interact with the mechanism that needs to be opened next (“state-advancing action”), by selecting the state-advancing action with probability 1 - *ϵ* and a random action otherwise (i.e., an *ϵ*-greedy strategy, Sutton et al. [30]). The exploration parameter *ϵ* ∈ [0, 1] controls the degree of randomness in the strategy. For *ϵ* = 1, the “smart” strategy collapses to the “random” strategy.

**Figure 3:**
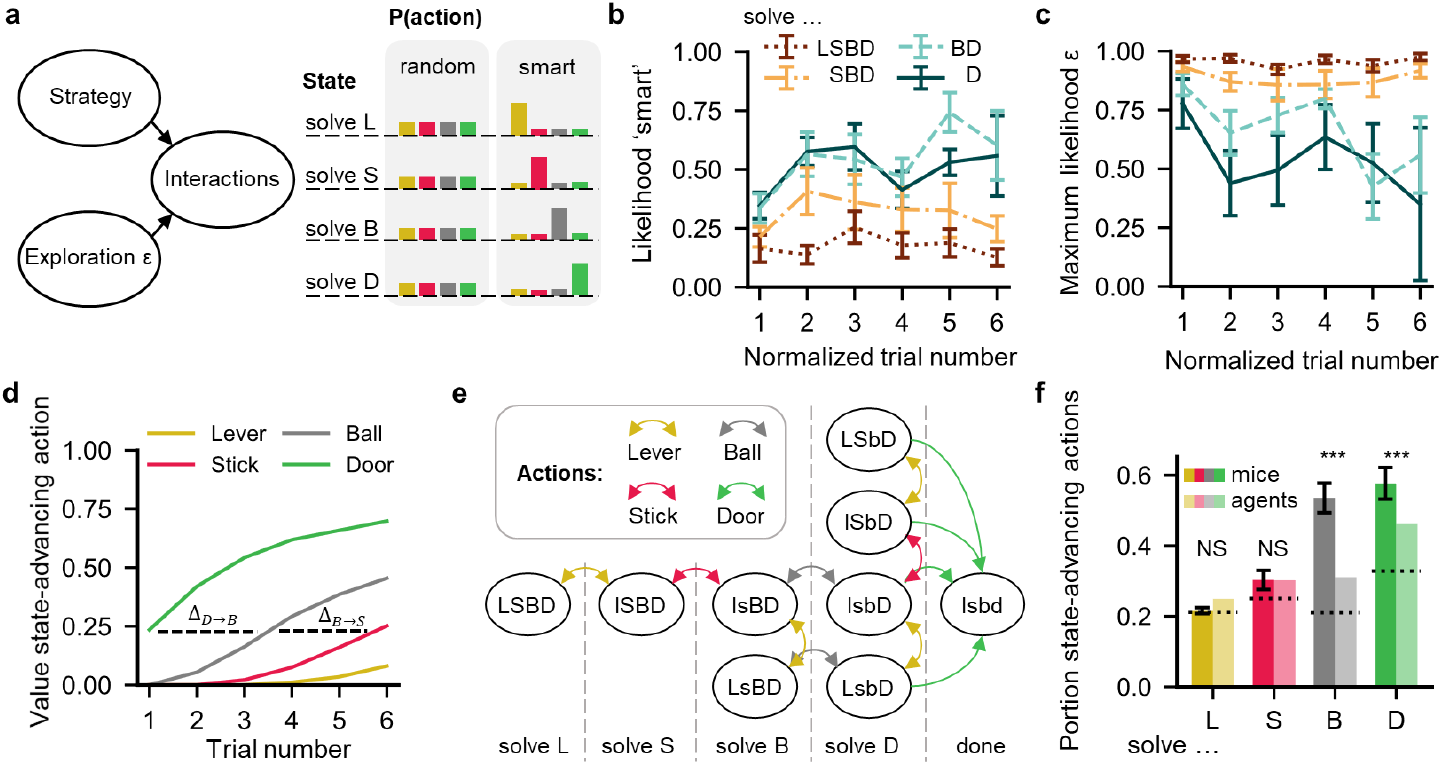
Solution strategies of mice solving the combined lockbox. **a**, Graphical model for the Bayesian inference and visualization of the behavioral policies for the two strategies. The states of the lockbox are named after which mechanism should be interacted with to advance the lockbox one step closer to its solution. **b, c** ‘smart’ model likelihoods and corresponding maximum likelihood epsilon values for the solved trials. The different curves correspond to different slices of data included in the inference (‘D’ indicating the ‘solve D’ state, ‘BD’ indicating both the ‘solve B’ and the ‘solve D’ state, etc.). Data represent the mean ±SEM, n = [12, 9, 8, 10, 6, 3] across trials 1 − 6. **d**, Average Q-values of state-advancing actions of simulated agents in a lockbox environment. Delays in reward learning between the door and ball action (Δ_*D*→*B*_) and between the ball and stick action (Δ_*B*→*S*_) are explicitly illustrated. **e**, Markov state diagram representing the combined lockbox. Closed and open mechanisms within a state are represented by an upper- and lowercase letter, respectively (e.g., “L” and “l”). Actions with nonzero probability of transitioning to a different state are color-coded, while ‘wrong’ actions, i.e., actions that do not yield state transitions, are omitted to increase readability. The Markov states are ordered to match the lockbox states from a and are separated by dashed lines. **f**, Average portion of state-advancing actions for mouse trials (left bars) and simulated agents (right bars). Error bars on the mouse trials indicate the standard error of the mean (n = [67, 59, 51, 50] for ‘L’, ‘S’, ‘B’, and ‘D’). The dotted lines represent the average mouse preferences for the four different mechanisms.

To infer which strategy the mice used, we calculated the likelihood of each strategy given the interactions performed by the mice. To ensure interaction data for all lockbox states, we only considered successful trials. When interactions with all mechanisms are taken into account, the random strategy is most consistent with the observed interaction data (Figure 3b, c, ‘LSBD’). However, based on reinforcement learning theory, we reasoned that learning could progress by backpropagating rewards to increasingly earlier stages of the task [30]. Thus, mice should initially realize that opening the door leads to the reward, later learn that the ball should be removed to be able to open the door, then learn to remove the stick, and lastly learn that the lever should be opened first. We illustrate this propagation of information about the underlying task structure across trials in a simulated lockbox environment (using temporal difference learning; Sutton et al. [30]; see Methods). In this model, the cumulative future value of opening each mechanism is updated by a temporal difference error, which compares the obtained reward to the difference between the values of the current and the next state. This leads to a delayed learning of earlier task stages (Figure 3d).

Following this reasoning, we repeated the inference analysis for increasingly late task stages. We find that for the last two stages, in which either only the door or only ball and door need to be solved, the smart strategy becomes increasingly likely across trials (Figure 3b, c; ‘BD’ and ‘D’). These findings are consistent with evidence that animals exhibit different types of backward learning processes from reward-proximal states [31, 32]. The reason why this emerging strategy is not reflected in the overall analysis is that the number of interactions in the early stages of the task greatly outnumbers the number of interactions in later stages (lockbox entirely closed: 2678 interactions, 1 open mechanism: 1090, 2 open: 431, 3 open: 238).

### A computational modeling to dissect learning

For a more detailed analysis of the behavior of the mice, we developed a computational model of the lock-box task using a Markov decision process (Figure 3e), which provides a baseline for random behavior and allows a gradual inclusion of different forms of learning. For a random baseline model, we included the preferences for the mechanisms of the mice, and matched the transition probabilities between the lockbox states to the mechanism-specific success rates in the mouse data. For a notion of successful and unsuccessful trials in the simulations, we defined an upper limit of 200 actions for the trial to be deemed successful. This limit was chosen such that the percentage of successful agent trials matches that of the mice (74% for mice, 75% for agents). The resulting average number of actions required to solve the task closely matches that of the mice (92 for mice, 98 for agents).

Consistent with delayed strategy learning, the behavior of the mice in the early task stages is described well by the random baseline model. The fractions of the state-advancing actions match those of the mice for the first two task stages (‘solve L’ and ‘solve S’ states, Figure 3f). Consistent with the Bayesian inference results, mouse behavior deviates from random behavior only for later task stages. State-advancing actions are overrepresented and differ significantly from the random baseline model (two-sample KS-test, *n*_*mice*_ = [67, 59, 51, 50], *n*_*agents*_ = 10, 000, ‘solve L’ p = 0.268, ‘solve S’ p = 0.098, ‘solve B’ p = 7 *×* 10^−17^, ‘solve D’ p=1 *×* 10^−6^).

### Motor learning dominates the rapid learning progress

Given that the overall learning progress is influenced by habituation, motor learning, and an emerging strategy based on task understanding, we aimed to assess the relative contributions of these three types of learning. To this end, we incrementally included the different types of learning in the computational model. As a baseline, we used the random model with state-transition probabilities matching the mechanism opening/closing probabilities of the mice in the first combined lockbox trial (“unskilled” agents, Figure 4a). To capture the effect of habituation, we mimicked the delayed engagement of the mice as a reduction in the upper limit of actions in the first trial before the trial was deemed a failure. To mimic motor learning, we increased the state transition probabilities to match the trial-specific success rates of the mice (Figure 2c). Finally, we added the development of a behavioral strategy as inferred by the statistical analysis (Figure 3b, c).

**Figure 4:**
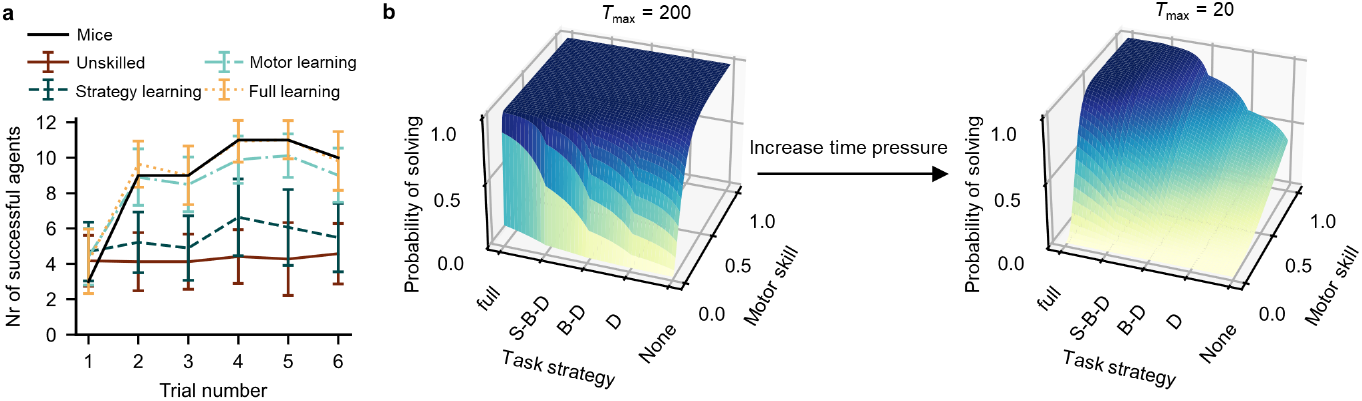
Learning occurs in both mechanism manipulation and understanding of the task structure. **a**, Number of successful mice/agents (out of a group of 12) across trials. Simulated agents have prior preferences based on the average preference of the mice. The “Full learning” agents contain both motor learning and the development of a strategy, while the “Unskilled” agents contain neither. The default success/failure threshold is set to 200 actions. Error bars represent standard deviations (n = 100). **b**, Probability of solving the (simulated) lockbox task as a function of mechanism manipulation skill, task understanding, and number of time steps (*T*_max_).

We find that habituation alone only has a modest effect on the number of successful trials. This resulted in a reduction of successful agents in the first trial from 4.17 (unskilled agents) to an average of 3.7 agents. The inclusion of motor learning, in contrast, generates a close match to the learning progress of the mice, which is only moderately improved in later trials by the additional inclusion of the emerging behavioral strategy (“Full learning” in Figure 4a). This indicates that the rapid learning observed in the mice is dominated by the increasing ability of the mice to manipulate the individual mechanisms. Nonetheless, the acquisition of motor skills alone cannot explain the full observed performance of the mice. It is the combination of both motor learning and the development of a cognitive strategy that closely matches the observed behavior.

### Motor learning is effective in the absence of time pressure

To understand which aspects of the task design favored motor learning over developing an understanding of the task, we used the computational model to systematically analyze the influence of motor skill and cognitive strategy on the probability of solving the lockbox trials (Figure 4b). Motor skill is modeled by increasing the probability of successfully opening/closing a mechanism when interacting with it (i.e., transition probabilities in the Markov decision process). For simplicity, all four mechanisms are assigned the same success probability. Cognitive strategy is modeled by decreasing the probability *ϵ* of random actions for the different mechanisms. Starting from fully random behavior, we reduce randomness sequentially for the door, ball, stick, and lever, mimicking the backpropagation of reward information as seen in reinforcement learning theory (see Methods). We also varied the time pressure by varying the upper limit of interactions before a trial was deemed a failure.

For the upper limit of 200 actions that matches the behavioral data best, we find that small changes in motor skill generate large performance gains, whereas task understanding causes only modest gains, at least for the initial poor motor skill. This is a result of the relative simplicity of the task, which can be solved by random behavior within a reasonable number of actions. With increasing time pressure, a non-random strategy becomes increasingly important (Figure 4b, *T*_max_ = 20), suggesting that an effective training paradigm for mice on the lockbox task should start with a period that favors skill learning, e.g., by single mechanism training, before exposing the mice to the full lockbox under increasing time pressure.

## Discussion

For mice, the lockbox is a novel sequential decision-making paradigm which provides a unique platform to investigate ethologically relevant learning behavior. Given the learning speed observed in conventional binary decision-making tasks [18, 19], the mice demonstrated remarkably fast learning by consistently solving the lockbox within a few trials. By combining pose tracking, Bayesian inference, and computational modeling, we were able to isolate distinct components of the learning process while allowing the mice to explore the cage freely. We showed how both motor learning and an emerging task strategy contributed to mice becoming increasingly capable of solving the task. The acquisition of motor skills occurred early and was the primary contributor to their learning process. Acquiring a strategy was slower and progressed in reverse order, with the mechanism closest to the reward learned first. This suggests that mice have acquired some knowledge about the task and switch from exploratory to exploitative behavior within the trial.

We were surprised by how effectively mice solved this complex task and that their development of manipulation skills appears to be the main contribution to their success. However, from an ecological perspective, a fixed, or even random strategy may initially be a very effective heuristic. Learning is beneficial only when the same problem setting occurs consistently over time, such that the effort of learning a task-specific behavior is amortized later [33, 34, 35]. If the problems encountered are highly variable, a random strategy may in fact be a good choice, albeit probably inferior to the more demanding variant of mental simulation. Instead, effort may be better spent on learning how to manipulate objects, granted that these are more likely to reappear at a later stage in their lifespan. Depending on the organism and type of task, this might furthermore come with lower costs than learning the task structure [36, 37, 38]. Indeed, there is accumulating evidence across species that higher-level cognitive processes are metabolically costly [39, 40, 41], while sensory-motor processes impose only minor additional costs [42]. Yet, as we describe in our simulations, random strategies are only effective in decision problems with a relatively small number of actions. If either the action sequence or the time pressure is too high, a solution strategy is necessary to consistently solve the problem and thus any additional costs of strategy learning must be overcome.

A potential inroad to a deeper understanding of how mice learn motor skills and abstract strategies for sequential problems would be to study how they generalize to new variants of the lockbox. If the increased performance was primarily due to motor learning or the identification of promising candidate actions, they should generalize to lockboxes composed of the same mechanisms in a different order. If they acquired an abstract understanding of sequential problems, their performance on entirely new sequential problems should improve with experience. An additional direction would be to vary the number of sequential steps in the lockbox, to probe the limits of strategy learning. At a certain point, increasing or decreasing task complexity may reveal a shift from structured strategies to reliance on motor skill and fixed or stochastic action patterns only, offering a way to disentangle flexible planning from habitual execution. Thus, the lockbox paradigm can provide unexplored pathways to studying the role between low-level motor learning and the acquisition of a task strategy, while providing clear anchor points for the analysis of both the behavior and its neuronal underpinnings.

## Methods

### Ethics statement

Animal experimentation was performed according to the guidelines of the German Animal Welfare Act and the Directive 2010/63/EU for the protection of animals used for scientific purposes. Maintenance of mice and all animal experimentation were approved by the Berlin State Authority (“Landesamt für Gesundheit und Soziales”, permit number: G0249/19). The experiment was pre-registered in the animal study registry (animalstudyregistry.org, doi: 10.17590/asr.0000237), comprising several substudies, alongside the analysis of lockboxes as enrichment, which is published in an accompanying paper [26].

### Experiments

#### Animals

12 female C57BL/6J mice were obtained from Charles River Laboratories (Sulzfeld, Germany). They were free of pathogens listed in the FELASA recommendation [43]. The mice arrived at the age of four weeks at the animal facility, where they were housed in filter top cages under specifically pathogen free barrier conditions. The mice were maintained under a room temperature of 22 ± 1 °C, a relative humidity of 55 ± 10%, and a 12/12-hour light/dark cycle. Lights turned on at 7 AM (summertime: 8 AM); 30 min before, a simulated sunrise started and smoothly increased the light intensity (Philips HF 3510, 100-240 vac, 50-60 Hz, Philips GmbH Market DACH, Hamburg, Germany). The mice were housed in groups of four in two Makrolon type III cages (39 × 23 × 15 cm) that were connected to each other by a gate. The floor of both cages was covered with fine wooden bedding material (JRS Lignocel FS14, spruce/fir, 2,5-4 mm). One of the cages contained a red triangular plastic house with two exits (long side: 14.5, short sides: 11.5, height: 5cm, TheMouseHouse, Tecniplast), a transparent handling tunnel (length: 11.5, outer diameter: 4cm, custom-built from GEHR PMMA XT® ACRYL, Mannheim, Germany), and three thin (cellulose, unbleached, layers, 20×20cm, Lohmann & Rauscher International GmbH & CO KG) as well as two thick (23×24,8cm folded, Essity ZZ Towel) paper tissues for nesting; the other cage was not enriched with any material and was considered as training cage. Food pellets (LASvendi, LAS QCDiet, Rod 16, autoclavable) and tap water were provided *ad libitum* in both cages. The animals were handled using a tunnel. In the first two weeks, they were habituated to tunnel handling by the animal technicians and experimenter, who were all female. After the mice had habituated to the animal facility for a week, radio frequency identification (RFID) transponders were subcutaneously implanted into the neck region using an application cannula (ISO-Compliant Transponder, Peddymark Limited, Redhill, Surrey, United Kingdom; cannula: 2.6 × 0.15 × 40 mm) under a short isoflurane anesthesia (induction: 4% isoflurane, maintenance: 1-2% isoflurane; carrier gas: 100% oxygen, flow: 1 liter/min; Isofluran CP 1ml/ml, CP-Pharma Handelsgesellschaft mbH, Burgdorf, Germany). Before the transponder implantation, 1 mg/kg meloxicam (Loxicom 0.5 mg/ml, Elanco Animal Health, Bad Homburg, Germany or Metacam 0.5 mg/ml, Boehringer Ingelheim, Ingelheim am Rhein, Germany) was orally administered. Besides the RFID transponders, color marking of the tail was used for rapid visual identification of the individuals (edding 750 paint marker, edding International GmbH, Ahrensburg, Germany). Before the lockbox training period, the mice were tested in the open field, elevated plus maze, free exploratory paradigm, grid exploratory paradigm, and calorimetry including presentation of a 2-step lockbox for another substudy [26].

#### Home cage set-up

The home cage of the mice consisted of two Makrolon type III cages connected to each other by a gate. The gate was made of three transparent tubes (11.5 × 4 cm, custom-built from GEHR PMMA XT® ACRYL, Mannheim, Germany) and two doors (custom-built, 3D printed, STL files can be found on Zenodo: http://www.doi.org/10.5281/zenodo.11143666). The doors could be opened and locked by the experimenter. Except during the training periods, the mice had access to both cages. During the training periods, the doors of the gate were closed and the mice could only individually enter the training cage. The cage lid of the training cage was modified for video monitoring and the grid was replaced with transparent acrylic glass in an area of 22 × 15 cm.

#### Video monitoring

Three cameras acA1920-40um (Basler AG, Ahrensburg, Germany; Kowa Lens LM25HC F1.4 f25mm, Kowa Optimed Deutschland GmbH, Düsseldorf, Germany) were used to record the mice during the training periods from the top, front, and side (1936 × 1216, 30 frames/s, AVI format). The cage was additionally illuminated with two infrared lights (Synergy 21 IR-Strahler 60W, ALLNET GmbH Computersysteme, Germering, Germany).

#### Lockbox designs

The lockboxes were designed using Tinkercad (www.tinkercad.com). The STL files were exported from Tinkercad and the G-code was generated with Cura SteamEngine 4.4.0. The lockboxes were printed using an Ultimaker 3 Extended and an Ultimaker S3 (0.4 mm nozzles). Different colored polylactic acid (PLA) was used as material. The lockbox consisted of a white platform, a yellow lever, a red stick, a gray ball, and a green sliding door. Information on 3D printing details and the STL files of the lockbox can be found on Zenodo [44].

#### Lockbox training

After the mice had been kept for four weeks in the animal facility, they were habituated to the food reward, i.e., oat flakes (kernige Haferflocken, Wurzener Nahrungsmittel GmbH, Wurzen, Germany). On three consecutive days, eight oat flakes were scattered on the floor of the training cage in the morning. On the following day, the lockbox training started. For this, the lockboxes were baited with an oat flake. There was one lockbox for each mouse to exclude odor cues from conspecifics. The lockbox training was carried out as follows: After all mice left the training cage, the doors of the gate were locked and the closed lockbox was placed into the training cage. Then the first mouse was allowed to enter the training cage and explore the lockbox for a maximum of 30 min. A training trial ended after the mouse opened the lockbox or after 30 min. At the end of the trial, the lockbox was removed from the training cage when the mouse no longer touched the lockbox with any paw or 1 min after the end of the trial. Finally, the doors of the gate were opened and the mouse could leave the training cage. Then another lockbox was placed into the training cage and the procedure was repeated for all mice over six trials. In all trials, the mice entered the training cage in a randomized order. In trial 1, the mice explored the closed lockbox for the first time. Between combined trial 1 and 2, the mice were trained on the single mechanisms of the lockbox (i.e., the lever, the stick, the ball, and the sliding door), over a period of 20 days, including a few days without lockbox training. There were 11 consecutive days of training on single mechanisms, followed by a 2-day break, 2 consecutive days of simultaneous presentation of all single mechanisms, and then another 5-day break. The single mechanisms were baited with half an oat flake. In combined trial 2, the open lockbox, baited with a full oat flake, was presented to the mice for 1 minute. After that, it was re-baited with a food reward and fully closed. The lockboxes were cleaned with ethanol 70 % after all mice completed their training trial.

### Tracking

#### Key point extraction in 2D

The positions of the mouse body parts and lockbox were tracked in each video using DeepLabCut (DLC, Mathis et al. [27], Nath et al. [28], version 2.2.1.1). Two individual trackers were trained to locate different key points, one for the lockbox (both ends of the lever, the head of the stick, the center of the ball, and the center of the sliding door), and one for the mouse (nose, both ears, base of the tail, and all paws). Among the full dataset of 3934 labeled images, 3554 (90%) were randomly assigned for training and 380 (10%) for testing both trackers. To extract the images from the videos and to train the networks, methods provided by DLC were used with default settings (see Suppl. Inf.).

#### Triangulation

The known 3-dimensional coordinates of the lockbox CAD model were used as the base coordinate system and were matched with the corresponding (automatically detected) 2-dimensional lockbox key points to generate a mapping from 3 videos *×* 2D observation space to 3D. For combined lockbox trials, we automatically identified frames in which the lockbox was closed, based on the distance between the tip of the lever and the head of the stick in both the front and the top perspectives. We used threshold values of *<* 610px horizontally and *<* 160px vertically in the front view, and *<* 383px horizontally and *<* 182px vertically in the top view. We retained up to 1000 frames of closed lockbox coordinates whose tracking likelihoods exceeded a threshold confidence (default: 0.8) and paired each triplet of 2D observations with its corresponding 3D position (from CAD drawings) for the triangulation step.

For the single mechanism trials we manually selected points on the lever lockbox from all three perspectives for each trial. We used the 4 corners of the base plate, the 4 corners of the elevated platform, the 4 corners of the lever support, and the two tips of the lever for the mapping. For the other single mechanism lockboxes (stick, ball, and door), we used the same manual key points from the lever lockbox of that trial (recorded on that same day), but applied a rotation around the z-axis on the 3-dimensional lockbox coordinates before constructing the mapping. The rotation to the 3D coordinates serves to correct for potential rotational displacements of the lockboxes by the experimenter.

To determine the rotation angles, we took the frames at 10%, 25%, 50%, and 75% of the top view videos of the stick, ball, and door lockbox, extracted a central region (75% of image dimensions), applied edge detection (gradient threshold = 70) and a probabilistic Hough line transform (vote threshold = 50, minimum line length = 200px, maximum gap = 40px) to find the right vertical plate edge (the edge most clearly visible in all frames). We averaged these angles (after ensuring that variance *≤* 0.05 rad^2^) and compared them with the orientation of the baseline plate, extracted from the manually selected key points. Finally, we applied this rotation correction to the initial 3D lockbox points.

To create the mapping, we used RANSAC to fit **x** = *H***r** + **x**^(0)^ by sampling *N −* 2 points per trial, requiring at least *N/*4 inliers with reprojection mean-squared error *≤* 100 px^2^ on a maximal of *N* + 10 iterations, where *N* is the number of frames, **r** are the coordinates in 3D, and **x** are the coordinates of the video triplets (6-dimensional: 2 *×* 3 perspectives). From the inlier set, we estimated the 6 *×* 3 matrix *H* and 6D vector **x**^(0)^, then computed their pseudo-inverse [*H*|**x**^(0)^]^+^ for back-projection and manifold-threshold filtering.

#### Manifold filtering

The 2D key points of the three cameras were concatenated into a 6D observation vector **z**(*t*). Before applying an (extended) Kalman filter to the data, we first removed any DLC detections that were not in the projection manifold: given **z**(*t*), we back-projected

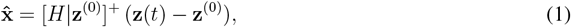

reprojected

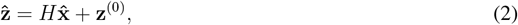

and discarded **z**(*t*) (i.e., set the confidence of the tracked key point to 0.01) if 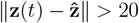.

#### Temporal filtering

(Extended) Kalman filters were used to refine the locations of the mouse and lockbox mechanisms in 3D. The underlying generative model maps the degrees of freedom of the mouse and the lockbox to the 6D coordinates of all key points obtained from the DeepLabCut tracker. We used a rigid-body model for the mouse and the lever, while the other mechanisms (ball, stick, and door) consisted of single points. The models were defined in terms of a global position **x**^cm^ (i.e., the “center of mass”) and orientation (pitch *θ*, yaw *ϕ*, and roll *ψ*) of the object and a set of angles describing the relative rotations between the rigid bodies in the model. The distances of the rigid bodies were chosen in line with a typical mouse, but not fine-tuned to the individual mice. The set of parameters for each model is listed in Extended Data Table 1, and the relative positions for the kinematic mouse and lever models are listed in Extended Data Table 2. Rotation matrices were used to define the orientations of the rigid parts in the model, which were implemented according to [45]. The order of the rotations around the axes was defined as y → z → x.

The skeleton motion kinematics and the individual mechanisms were assumed to behave according to a constant acceleration model:

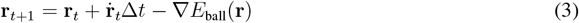

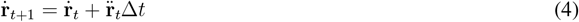

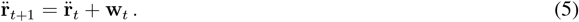

Here, **r** = *{***x**^cm^, *θ, ϕ, ψ}* denotes the parameters of the model, Δ*t* is the discretization time step, and **w** ∼ *𝒩* (0, **Q**) is the process noise, drawn from an *N* -dimensional Gaussian. **Q** denotes a diagonal covariance matrix that characterizes the amount of noise in dynamics that can be accounted for by the observations **z**. *E*_ball_(**r**) is an energy function and the term −∇*E*_ball_(**r**) biases the state estimate towards regions of small energy. This was used for the ball to incorporate prior knowledge. We furthermore added angular dampening to the orientation parameters of the mouse model to avoid the model from spinning out of control. The exact values can be found in Extended Data Table 1.

Different lockbox mechanisms have different degrees of freedom. The lever is characterized by a single tilting angle ranging from 0 to *π/*2. The stick and the sliding door are described by a 1D variable indicating how far it has been shifted from an arbitrary zero position. The ball is free to move in 3D space, but we included an energy function

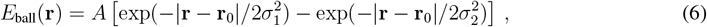

which favors finding it at its original position **r** in the horizontal plane, unless it had previously been removed.

Finally, the 3D coordinates **x**_*i,t*_ of the different key points are converted into 6D tracks *z*_*i,t*_ using the linear transformation defined in the triangulation section,

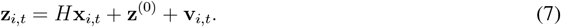

The observation noise **v**_*i,t*_ ∼ 𝒩 (0, **R**_*i,t*_) accounts for errors in the key point observations. The scale of this noise was defined by the diagonal matrix **R**_*i,t*_. We included the confidence *c ∈* [0, 1] of the DLC tracker by scaling the observation noise for key points with a low confidence:

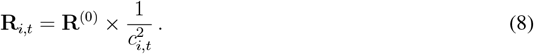

This scaling is applied individually for each tracked key point and camera. The effect of this rescaling is to ignore unreliable observations and infer the respective keypoint from either the other camera views or from the assumed continuity in time. Such unreliable observations are frequent due to occlusions.

### Behavioral analyses

#### Interaction analysis

Interactions with the different mechanisms of the lockbox were detected by defining 3D bounding boxes around each lockbox part, chosen manually. Each video frame was classified as an interaction of the mouse with a given lockbox part when the mouse’s nose was in the associated bounding box. These Boolean sequences were smoothed with a 30-frame weighted moving-average filter (Gaussian-like weights, threshold = 0.3) to remove noise. Any frame whose KF covariance diagonal exceeded 10 mm^2^ in *x, y*, or *z* was ignored in the analysis to remove frames in which the mouse is likely not in the observable front part of the cage. The smoothed sequences were used to calculate interaction times with the lockbox mechanisms. To assure that the interactions with the mechanisms were labeled in the correct lockbox states, we manually annotated the lockbox states in each trial.

#### Bayesian analysis of behavioral strategies

For a more fine-grained quantitative analysis of the strategic behavior of the mice, we performed a Bayesian analysis that tested whether the interaction pattern of the mice was consistent with two different strategies. We segmented the behavior into discrete interactions with the four lockbox mechanisms (lever - 1, stick - 2, ball - 3, door - 4). An interaction was defined as an uninterrupted sequence of interaction-detected frames with a single lockbox mechanism. To limit the number of ‘unintended’ interactions in our data (e.g., merely walking over the mechanisms, or false positive frames), we only consider uninterrupted interactions of at least 1.5 s, such that it approximately corresponds to the median overall interaction time across all trials (1.47 s). The model also considers four different lockbox states, which correspond to which mechanism of the lockbox had to be opened next to proceed (“solve L”- 1, “solve S” - 2, “solve B” - 3, “solve D” - 4). For our Bayesian analysis, we only considered trials in which the animal solved the task, such that interaction data for all lockbox states are available.

The strategy model assigns a conditional probability *π*(*i*|*j*) = *π*(interaction *i*|lockbox state *j*) to interact with each mechanism of the lockbox, given the overall state of the lockbox. We compared two different strategy models. The first strategy was random behavior, in which the mouse interacted with the different lockbox mechanisms with probability equal to its overall empirical probability, irrespective of the lockbox state:

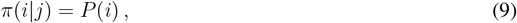

where *P* (*i*) denotes the prior preference distribution of the mouse, which is determined by calculating the fraction of interactions per mechanism across all combined trials for that mouse. The second strategy describes goal-directed behavior with interactions biased toward the lockbox mechanism that need to be solved next to progress the task:

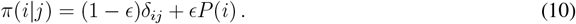

Here, *δ*_*ij*_ denotes the Kronecker symbol, which is 1 if *i* = *j* and zero otherwise. The parameter *ϵ* ∈ [0, 1] controls the amount of randomness (or exploratory behavior) of the mouse. For *ϵ* = 1, the behavior reverts to the above random strategy, for *ϵ* = 0, the mouse interacts exclusively with the lockbox mechanism that needs to be opened next.

Assuming that the interactions are statistically independent samples from the distribution *π*_*ij*_, this model allows to compute the likelihood of the randomness parameter *E* for any sequence (*I*_*m*_, *s*_*m*_) of interactions *I*_*m*_ and states *s*_*m*_:

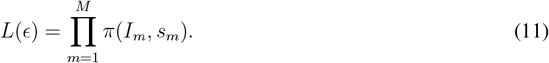

The model also allows to perform model comparison, i.e., the likelihood of the two different strategies given the observed state-interaction sequence. We evaluated *L(ϵ)* on a uniform grid of *ϵ* values (step = 0.025) and compared the results of the random and smart models to select the strategy with the maximum likelihood.

### Simulations

#### Environment

All simulations were performed using a Markov decision process (MDP) with 9 states: eight unsolved lockbox configurations (labeled as “LSBD”, “lSBD”, “lsBD”, “LsBD”, “lsbD”, “lSbD”, “LsbD”, “LSbD”, where the upper and lowercase letters represent a closed and opened mechanism, respectively) and a terminal absorbing state (“lsbd”) corresponding to solving all the mechanisms, illustrated in Figure 3e. At each time step, agents selected one of the four actions *A* = {lever, stick, ball, door}. The transition probabilities of the MDP were parameterized by the opening and closing success rates per mechanism *p*_*a*_ (one for each of the four actions *a ∈ A*). The simulations ended when reaching the exit state “lsbd”.

#### Static agents

We incorporated different modalities of learning as follows. The development of motor skills was simulated by increasing the transition probabilities of the MDP for all agents, whilst keeping the policy of the agents unchanged. A cognitive strategy was incorporated by using *E*-greedy agents with *E* values matching those inferred from the mouse data (see previous section). Epsilon values were assigned per state of the lockbox, meaning that an agent for example could in principle adopt a random strategy for the “solve L” state, while being goal-directed for the “solve D” state. For the comparison between agents and mice, the probability of an agent adopting an *E*-greedy strategy in a given lockbox state and trial equaled to the portion of mice solving the lockbox of that trial.

#### Learning agents

In our simulations to demonstrate the effect of delayed reward learning, we used an *E*-greedy policy combined with the SARSA learning rule [30]:

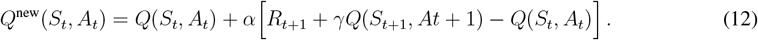

The agents were initialized with zeroes for the Q-values, a learning rate of *α* = 0.5, exploration parameter *E* = 0.3, and discount factor *γ* = 0.99. Upon solving the entire lockbox, the agent received a reward of *R* = 1. All other states were not rewarded. Since this simulation merely served the purpose of demonstrating the backpropagation of reward information across trials, we assumed 100% motor skill, i.e., the transition probabilities for the correct actions were 1. To obtain smooth learning curves, we used the average Q-values of 100 agents.

#### Solve probability versus task strategy & motor skill

To simulate the influence of task strategy and motor skill on the probability of solving the lockbox task within *N* steps, we used the absorption property of Markov chains. Given a specific fixed policy (task strategy) and set of state-transition probabilities (motor skill), we defined the lockbox task as a Markov chain and computed the cumulative absorption probability after *N* steps:

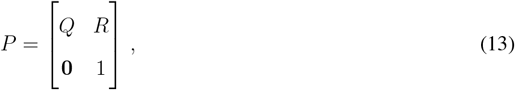

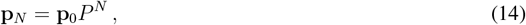

where *Q* is the transient state matrix, *R* is a submatrix for transitions from a transient to the absorbing state, and **p**_0_ is a vector representing the initial state of the agent (which is always at “LSBD”). Since the agents cannot escape the “lsbd” or “task solved” state after entering it (as it is an absorbing state), the last value of **p**_*N*_ represents the probability of the agent solving the task within *N* steps.

The absorption probabilities are computed for a grid of agents with one dimension representing their motor skills (with a step size of 1/100), and the other dimension representing task strategy. For simplicity, all four mechanisms are assigned the same motor skill, i.e., state transition probability. Cognitive strategy is modeled by decreasing the probability *ϵ* of random actions for the different mechanisms. Starting from fully random behavior, we reduce randomness sequentially for the door, ball, stick, and lever in favor of selecting the state-advancing action, mimicking the backpropagation of reward information as seen in reinforcement learning theory. The labels “None”, “D”, “B-D”, “S-B-D”, and “full’ represent which mechanisms of the lockbox are already learned. For already learned states (e.g., ball and door from the label ‘B-D’ until the label ‘S-B-D’), we assume optimal decision making (always select the state-advancing action), while for to-be learned state (stick in this example), an *E*-greedy policy is used. The agents adopt a random strategy for the lever in the ‘B-D’ → ‘S-B-D’ range, since this step will only be learned from the ‘S-B-D’ range onward.

### Statistical information

All regression-based statistical tests are computed by performing linear regression using ‘linregress’ from the Scipy package in Python [46]. This function uses a two-sided Wald Test whose null hypothesis is that the slope is zero.

Due to errors in some recordings, data from a few trials are missing.

- The number of used samples for the linear regression for combined lockbox interaction times (Fig. 2d) from trial 1 to 6 are: [12, 12, 10, 10, 11, 12] when including all trials (black curve of the figure). The number of used samples for the solved and unsolved trials are [12, 9, 8, 10, 6, 3] and [9, 6, 2, 1, 1, 0], respectively. Note that for the latter two the decaying number of samples naturally occurs due to the nature of what is being tested: not all mice solve the task on every trial, and no mouse does not solve the task at all.
- The number of used samples for the linear regression for combined lockbox preferences (Fig. 2e) are: [12, 12, 10, 10, 11, 12].

The error bars of the single mechanism interaction times (Fig. 2b) consists of [12, 11, 12, 11, 12, 10, 11, 8, 9, 12, 12], [12, 11, 12, 10, 12, 10, 10, 8, 9, 11, 12], [11, 11, 12, 11, 12, 10, 9, 8, 9, 12, 12], and [11, 11, 12, 10, 12, 10, 10, 9, 9, 12, 12] data points for the lever, stick, ball, and door, respectively.

For the comparison of portion state-advancing actions between mice and random agents (Fig. 3f) we used a two-sample Kolmogorov-Smirnov test. The sample sizes for the mouse data were n = [67, 59, 51, 50] for the ‘solve L’, ‘solve S’, ‘solve B’, and ‘solve D’ states, and the sample size for the random agents was 10,000. We used the ‘ks-2samp’ function from Scipy.

## Data availability

The data presented here are available from the corresponding author on reasonable request.

## Computer code

The customized processing pipeline will be made publicly available upon publication.

## Acknowledgements

Funded by the German Research Foundation (DFG) under Germany’s Excellence Strategy – EXC 2002/1 “Science of Intelligence” – project number 390523135. H.S. performed parts of this work at the Okinawa Institute of Science and Technology (Japan) within its Theoretical Scientist Visiting Program. The authors thank Patrik Reiske and Leonie Pawlak for their help with manual data annotation and Clara Bekemeier for assisting in conducting the experiments.

## Author contributions

K.H. performed the experiments. K.H., N.A. and L.L. designed the 3D lockboxes. M.N.B. and S.T. performed the behavioral analyses. M.N.B. ran the Bayesian inference and simulations. M.N.B., N.A. and S.T. implemented the data processing pipeline. S.M. assisted in training the 2D tracker. M.N.B., S.T., N.A. and H.S. wrote the manuscript, and all authors contributed to revisions. O.H., L.L., C.T-R., and H.S. conceived the project.

## Footnotes

1 https://github.com/RefinementReferenceCenter/MouseLockBox/blob/main/Door_syste/Door_system.mp4

