## Supplementary Information & Extended Data for "Mechanical problem solving in mice"

#### 2D tracking

The dataset used to develop the trackers for mice and lockbox consists of a total of 79 videos showing the two different lockbox training conditions, i.e. training on single mechanisms of the lockbox and on the complete lockbox, and recorded from the three different perspectives (front, top, and side). From those 79 video recordings, 3934 images were extracted using the frame extraction method provided by DeepLabCut (DLC, version 2.2.1.1), keeping the default settings, that is, automatic extraction mode, kmeans algorithm for frame selection, no frame cropping, and a clustering step of 1 to include each frame for clustering. Among the full dataset, 3554 images (90%) were randomly assigned for training and 380 (10%) for testing both trackers. The composition ratio of the images of the two lockbox training conditions and the three perspectives, respectively, was maintained in the training dataset, while for the testing dataset the proportion of images for each case was evenly distributed. The detailed composition of the entire dataset as well as the training and testing sets is listed in Table 1 and Table 2. Each image was manually annotated with various keypoints on the mouse, such as nose, ears, paws, and tail, and on the lockbox (lever, stick, ball, and sliding door) using DLC's interactive graphical user interface as labeling tool.

| | Front | Top | Side | Total (%) |
|---|---|---|---|---|
| Single mechanisms | 380 | 490 | 327 | 1197 (30%) |
| Lockbox | 780 | 1117 | 840 | 2737 (70%) |
| Total (%) | 1160 (30%) | 1607 (40%) | 1167 (30%) | 3934 |

**Table 1:** The numbers of images taken of the two lockbox training conditions and the three different perspectives, respectively, used in the full dataset.

| | Single mechanisms | Lockbox | Front | Top | Side | Total (%) |
|---|---|---|---|---|---|---|
| Training set | 1007 | 2547 | 1030 | 1487 | 1037 | 3554 (90%) |
| Testing set | 190 | 190 | 130 | 120 | 130 | 380 (10%) |

**Table 2:** The numbers of images taken of the two lockbox training conditions and the three camera perspectives, respectively, used for training and testing sets.

Two individual trackers were developed to localize different keypoints on the lockbox and the mouse, respectively. The Lockbox Tracker and the Mouse Tracker were trained and tested independently, both following the same training and testing process and using the aforementioned image sets. The network configurations of each model were set to the default settings of DeepLabCut, employing the ResNet-50, a deep convolutional neural network architecture, pre-trained on the ImageNet dataset for pose estimation. The training and testing of the trackers was performed on a Linux system with 183-Ubuntu SMP kernel version and NVIDIA Quadro RTX 4000 GPU 8GB VRAM. Each training session had 1,030,000 training iterations, a batch size of 1, and a learning rate of $5 \times 10^{-3}$, retaining the multi-step learning rate decay as defined by DLC. The maximum input size of images was changed from 1500 by default to 2500 to allow larger frame sizes during training.

To evaluate the performance of the trackers, the accuracy, sensitivity, and precision analyses were performed and the pixel error between label and prediction was computed according to following equations:

$$\text{Accuracy} = \frac{true\,positive + true\,negative}{labels + predictions} \tag{1}$$

$$\text{Precision} = \frac{true\,positive}{predictions} \tag{2}$$

$$\text{Sensitivity} = \frac{true\,positive}{labels} \tag{3}$$

$$\text{Pixel Error} = \sqrt{(prediction - label)^2} \tag{4}$$

where $true\,positive$ and $true\,negative$ are the numbers of times, the trackers correctly predicted that a keypoint was present and not present, respectively. A point was considered correctly predicted with a tracker confidence $\geq 0.6$. The pixel error describes the distance between the predicted point and the labeled point for one keypoint in pixels. Note, that the above equations refer to the tracker predictions for one image and one keypoint. The overall performances of the trackers were calculated as the averages of all images and the number of keypoints. The results of the evaluation can be found in Table 3. The Lockbox Tracker performs slightly better than the Mouse Tracker, especially in terms of precision. This is probably due to the fact that the lockbox was always placed in the same position, whereas the mouse could move freely in the cage. As a result, the possible coordinates of the lockbox parts were limited in space and vary less, even if they were moved.

| Tracker | Mean Accuracy | Mean Precision | Mean Sensitivity | Mean Pixel Error |
|---|---|---|---|---|
| Lockbox | 87.94% | 96.68% | 84.93% | 27.68 px |
| Mouse | 84.95% | 89.83% | 67.92% | 17.07 px |

**Table 3:** Overall performance of the Mouse Tracker and the Lockbox Tracker.

#### Comparison between annotated labels and pipeline

To benchmark the accuracy of our data analysis pipeline, we compared the results of the tracking pipeline with manual annotations for two trials on a frame-to-frame basis. The two trials have a total duration of 22 minutes and 10 seconds. For each trial, two independent annotators marked the interactions between the mouse and the lockbox as well as the changes in the state of the lockbox mechanisms throughout the video footage. The automated pipeline results and the manual annotations are highly correlated (Figure 1). A more extensive comparison can be found in our dataset publication of the experiments.

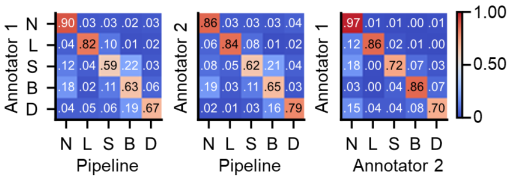


**Figure 1:** Confusion matrices comparing the detected interactions of the pipeline with two human annotators. The labels "L", "S", "B", and "D" represent interactions with the lever, stick, ball, and door, respectively. The "N" label represents the inactive time of the mouse.

#### Disengagement of mouse 68

We observe that one of the mice (ID=68) exhibited good initial performance in combined lockbox training. Specifically, in the first trial, it solved three mechanisms; in the second trial, it solved the complete lockbox; and in the third trial, it only solved one mechanism. However, it did not solve any mechanism in the fourth and fifth trials and solved only the first mechanism in the sixth trial. Figure 2 shows that the disengaged mouse started to disengage from the task after the third trial, and its engagement (ratio of time spent interacting with the lockbox over the total time) was lower than the engagement of the other 11 mice in the last three trials.

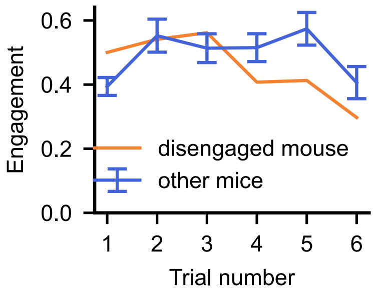


**Figure 2:** Engagement (ratio of time spent interacting with the lockbox over the total time) of the disengaged mouse compare to the other 11 mice. The error bars represent the standard error of the mean.

### Extended Data

| Object | Pose parameters | Description | Dampening $\tau$ (rotation), (velocity), (acceleration) |
|---|---|---|---|
| Mouse | $\mathbf{x}^{\text{cm}}$ | Position of the “center of mass” (base of the head) | - |
| | $\theta_1, \phi_1, \psi_1$ | Head rigid body pitch, yaw, and roll | (1, -, 0.5), (0.5, 0.5, 0.5), (0.1, 0.1, 0.1) |
| | $\theta_2, \phi_2$ | Neck rigid body pitch and yaw | (1, 1), (0.5, 0.5), (0.1, 0.1) |
| | $\theta_3, \phi_3$ | Back rigid body pitch and yaw | (1, 1), (0.5, 0.5), (0.1, 0.1) |
| Lever | $\mathbf{x}^{\text{cm}}$ | Position of the “center of mass”(hinge of the lever) | - |
| | $\theta$ | Lever rigid body pitch | - |
| Stick | $\mathbf{x}^{\text{cm}}$ | Position of the head of the stick | - |
| Ball | $\mathbf{x}^{\text{cm}}$ | Position of the center of the ball | - |
| Door | $\mathbf{x}^{\text{cm}}$ | Position of the center of the door | - |

**Extended Data Table 1:** Pose parameters for all tracked objects. The dampening $\tau$ for the angular parameters is implemented as an additional term $-\Delta t/\tau$ for their respective parameters.

| Object | Key point | Relation to global coordinate |
|---|---|---|
| Mouse | base of head ($\mathbf{x}^{cm}$) | $\mathbf{x}^{\text{cm}}$ |
| | nose ($\mathbf{x}^{\text{nose}}$) | $\mathbf{x}^{\text{cm}} + (\mathbf{R}_{\theta_1}\mathbf{R}_{\phi_1}\mathbf{R}_{\psi_1})^T[18.4\ \ 0\ \ -18.4]^T$ |
| | ear_left ($\mathbf{x}^{\text{ear left}}$) | $\mathbf{x}^{\text{cm}} + (\mathbf{R}_{\theta_1}\mathbf{R}_{\phi_1}\mathbf{R}_{\psi_1})^T[0\ \ 9\ \ 0]^T$ |
| | ear_right ($\mathbf{x}^{\text{ear right}}$) | $\mathbf{x}^{\text{cm}} + (\mathbf{R}_{\theta_1}\mathbf{R}_{\phi_1}\mathbf{R}_{\psi_1})^T[0\ \ -9\ \ 0]^T$ |
| | back ($\mathbf{x}^{\text{back}}$) | $\mathbf{x}^{\text{cm}} + (\mathbf{R}_{\theta_2}\mathbf{R}_{\phi_2})^T(\mathbf{R}_{\theta_1}\mathbf{R}_{\phi_1}\mathbf{R}_{\psi_1})^T[-24.6\ \ 0\ \ 4.3]^T$ |
| | tail_base ($\mathbf{x}^{\text{tail}}$) | $\mathbf{x}^{\text{back}} + (\mathbf{R}_{\theta_3}\mathbf{R}_{\phi_3})^T(\mathbf{R}_{\theta_2}\mathbf{R}_{\phi_2})^T(\mathbf{R}_{\theta_1}\mathbf{R}_{\phi_1}\mathbf{R}_{\psi_1})^T[-31.8\ \ 0\ \ -31.8]^T$ |
| Lever | hinge ($\mathbf{x}^{\text{cm}}$) | $\mathbf{x}^{\text{cm}}$ |
| | lever_tip ($\mathbf{x}^{\text{lever\_tip}}$) | $\mathbf{x}^{\text{cm}} + \mathbf{R}_{\theta}^T[65\ \ 0\ \ 0]$ |
| | upper_lever tip ($\mathbf{x}^{\text{upper\_lever\_tip}}$) | $\mathbf{x}^{\text{cm}} + \mathbf{R}_{\theta}^T[0\ \ 0\ \ 65]$ |

**Extended Data Table 2:** Mappings to find global key point locations based on the kinematic model, local rigid body distances, and rotation parameters (see Extended Data Table 1). The rotation matrices $\mathbf{R}_\theta$, $\mathbf{R}_\phi$, and $\mathbf{R}_\psi$ correspond to the pitch, yaw, and roll, respectively.
